# A highly conserved complete accessory *Escherichia coli* type III secretion system 2 is widespread in bloodstream isolates of the ST69 lineage

**DOI:** 10.1101/396804

**Authors:** S. Fox, C. Goswami, M. Holden, J.P.R. Connolly, A. Roe, M. Connor, A. Leanord, T.J. Evans

## Abstract

Bacterial type III secretion systems (T3SS) play an important role in pathogenesis of Gram-negative infections. Enteropathogenic and enterohemorrhagic *Escherichia coli* contain a well-defined T3SS but in addition a second T3SS termed *E. coli* T3SS 2 (ETT2) has been described in a number of strains of *E. coli.* The majority of *E. coli* contain elements of a genetic locus encoding ETT2, but which has undergone significant mutational attrition rendering it without predicted function. Only a very few strains have been reported to contain an intact ETT2 locus. To investigate the occurrence of the ETT2 locus in strains of human pathogenic *E. coli*, we carried out genomic sequencing of 162 isolates obtained from patient blood cultures in Scotland. We found that all 26 ST69 isolates from this collection contained an intact ETT2 together with an associated *eip* locus which encodes putative secreted ETT2 effectors as well as *eilA*, a gene encoding a putative transcriptional regulator of ETT2 associated genes. Using a reporter gene for *eilA* activation, we defined conditions under which this gene was differentially activated. However, comparison of secreted proteins from ST69 strains under high and low *eilA* activation failed to identify any ETT2 secreted substrates. The conservation of the genes encoding ETT2 in human pathogenic ST69 strains strongly suggests it has functional importance in infection, although its exact functional role remains obscure.

**Importance:** One of the commonest bacteria causing bloodstream infections in humans is *Escherichia coli*, which has a significant morbidity and mortality. Better understating of the mechanisms by which this microbe can invade blood could lead to more effective prevention and treatment. One mechanism by which some strains cause disease is by elaboration of a specialized secretion system, the type III secretion system (T3SS), encoded by the locus of enterocyte effacement (LEE). In addition to this well-defined T3SS, a second T3SS has been found in some *E. coli* strains termed *E. coli* type III secretion system 2 (ETT2). Most strains carry elements of the ETT2 locus, but with significant mutational attrition rendering it functionless. The significance of our work is that we have discovered that human bloodstream isolates of *E. coli* of sequence type 69 contain a fully intact ETT2 and associated genes, strongly suggesting its functional importance in human infection.

## Introduction

Pathogenic bacteria possess a number of different secretion systems that facilitate host infection as well as interbacterial competition(1). One of these is the type III secretion system (T3SS), which is found in a number of different Gram-negative pathogens and is key to the ability of these microbes to cause disease(2-4). Broadly, T3SS comprise two elements: a highly conserved multiprotein structural complex that forms the conduit between the bacterial and the host cell; and various effector proteins that are translocated through this channel. Genes encoding the T3SS channel, or needle complex, are contained within pathogenicity islands comprised of a single cluster of genes(5). Genes encoding effectors are more widely spread within the genome and vary greatly between different bacterial species.

Certain strains of *Escherichia coli* possess a well-defined T3SS, notably enteropathogenic *E. coli* (EPEC) and enterohaemorrhagic *E coli* (EHEC). This T3SS is encoded on the locus of enterocyte effacement (LEE) and in concert with its secreted effectors, produces the characteristic attaching and effacing lesions that mediate close attachment of the pathogen with the intestinal epithelial wall(6). Whole genome sequencing of strains of EHEC revealed the presence of a putative additional T3SS(7, 8), which has been termed *E. coli* T3SS 2 (ETT2). Further studies attempted to delineate the frequency with which this ETT2 locus was found in different *E. coli* strains(9-11). However, a further study by Ren et al(12) showed that the ETT2 locus was present in many lineages of *E.coli,* but had undergone extensive mutational attrition. The phylogenetic analysis showed that ETT2 was absent in what is thought to be the oldest phylogroup of *E.coli*, B2(13, 14), which contains many uropathogenic *E. coli,* but had been acquired by the divergence of the next oldest phylogroup, D. Analysis showed multiple inactivating mutations were present within the locus, which would render the T3SS functionless, including the ETT2 locus in the EHEC O157 strains in which it was originally described. However, a complete and potentially fully functional ETT2 was found in the enteroadhesive *E coli* O42 (EAEC O42) strain; other *E. coli* strains analysed either had no ETT2 locus, or it had undergone extensive deletion and/or mutational inactivation. Ren et al also showed that *E. coli* strains with the most intact ETT2 locus also carried an additional T3SS-like island adjacent to the *selC* tRNA gene, the *eip* locus, which encoded homologues of translocated proteins from the Salmonella pathogenicity island I (Spi-1) T3SS, as well as genes encoding a transcriptional regulator (*eilA*), a chaperone (*eicA*) and an outer membrane invasion/intimin-like protein (*eaeX*)(12, 15).

Functional effects of ETT2 remain unclear. Mutational analysis of the ETT2 cluster in an avian pathogenic *E. coli* showed it had reduced virulence, even though the cluster had undergone mutational attrition and could not encode a functional T3SS, suggesting potential alternative roles in pathogenesis(16). Other studies have also suggested a role for proteins encoded in the ETT2 in virulence of avian pathogenic *E. coli* and K1 strains(17-19). A recent study examined the role of the putative transcriptional regulator gene *eilA* at the *selC* locus in EAEC strain O42(15). This demonstrated that *eilA* was responsible for regulating transcription of genes within the *selC* locus, as well as *eivF* and *eivA* within the ETT2 locus. Mutants lacking *eilA* were less adherent to epithelial cells and had reduced biofilm formation; this phenotype was also observed for mutants in the *eaeX* gene which encodes the invasin/intimin homologue. This suggested important functional roles of the *selC* and ETT2 loci in pathogenesis of this strain of *E. coli*.

Hitherto, there is no evidence of intact ETT2 in human pathogenic strains of *E. coli* other than a few strains of EAEC. However, given the findings described above, we hypothesised that ETT2 might be of importance in human infections caused by *E. coli* phylogroups other than B2. We have studied 162 isolates of *E. coli* isolated from bacteremic patients in Scotland from 2013 and 2015, which we have subjected to whole genome sequencing. Within this group, we identified 26 strains of *E. coli* sequence type (ST) 69, of phylogroup D, which were largely derived from community-acquired sources. Virtually all of these strains had a completely intact ETT2 and *selC* operon, with no inactivating mutations. Similarly, intact ETT2/*selC* operons were also found in some minor ST types in our collection. The *eilA* transcriptional regulator was functional in these strains. Our results show that an intact ETT2 locus is widely present in human pathogenic *E. coli* ST69 strains, suggesting a functional role for this cryptic T3SS in human disease.

## Results

We have performed whole genome sequencing and analysis of 162 isolates of *Escherichia coli* obtained from blood cultures of patients within Scotland in 2013 and 2015(20). Sequence comparisons with other isolates of *E. coli* showed that strains belonging to ST69 contained an intact ETT2 operon. The gene content of this operon from one of these ST69 strains, ST69 1_9, was compared to the complete ETT2 found in enteroadhesive *E. coli* strain 042 (EAEC 042) and the degenerate ETT2 found in *E. coli* O157:H7 Sakai (Figure 1). An intact ETT2 operon in this ST69 strain was found in the ∼30 kb region spanning the *yqeG* gene and the tRNA gene *gluU* with over 98% identity to the ETT2 operon in EAEC 042. Importantly, this operon did not contain any of the inactivating mutations found in the *E. coli* O157:H7 Sakai strain.

**Figure 1.**
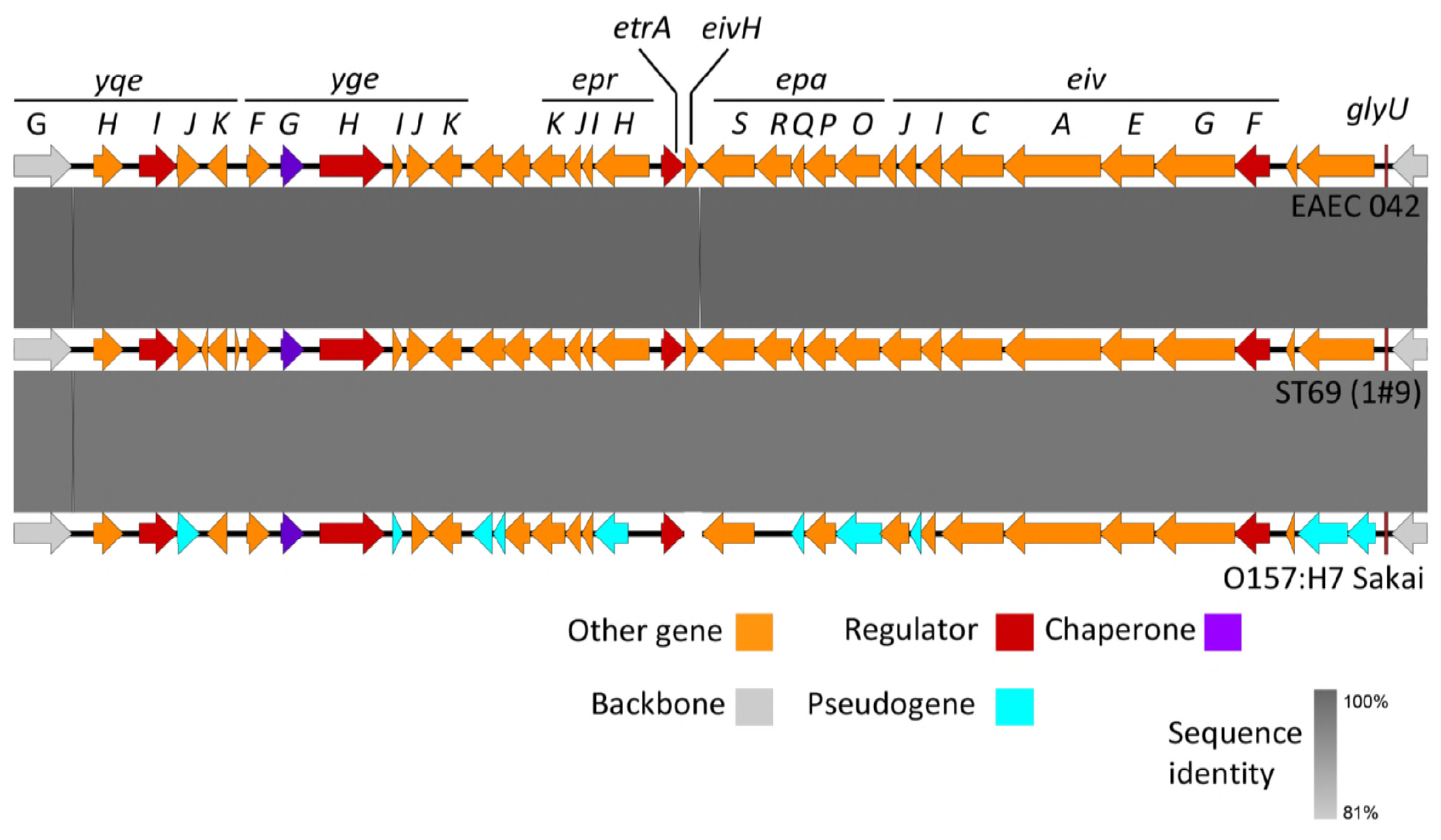
Comparisons of the ETT2 operon between EAEC 042, ST69 (1#9) and O157:H7 Sakai. Degree of identity is shown by the level of grey shading as indicated. Genes are colour coded according to putative function as shown.

We extended this analysis to compare all of the ST69 strains in our collection over this region. Of 26 ST69 genomes sequenced, 24 were assembled in one contig covering this region, shown compared to each other in Figure 2. In all these assemblies, there was a greater than 95% identity between the sequences. Two strains appeared to lack the extreme left-hand end of the complete ETT2 operon (ECO#35 and EC1#2), and two strains had a stop codon in the *epaO* gene at the same site as noted for *E. coli* O157:H7 Sakai (EC1#70 and ECO1#18). *epaO* is homologous to the *Salmonella typhimurium* Type III secretion system gene, *spaO*, which encodes a protein that forms part of the cytoplasmic sorting platform essential for energizing and sorting substrates for delivery to the needle complex(21). *spaO* is essential for type III secretion in *S. typhimurium(22)*. Recent work has shown that *spaO* produces two protein products by tandem translation: a full-length protein and a shorter C terminal portion that is translated from an internal ribosome binding site and alternative initiator codon(23). Both are needed for functionality of the type III secretion system in *Salmonella typhimurium*, so the loss of the full-length product of *epaO* will likely also render the ETT2 non-functional.

**Figure 2.**
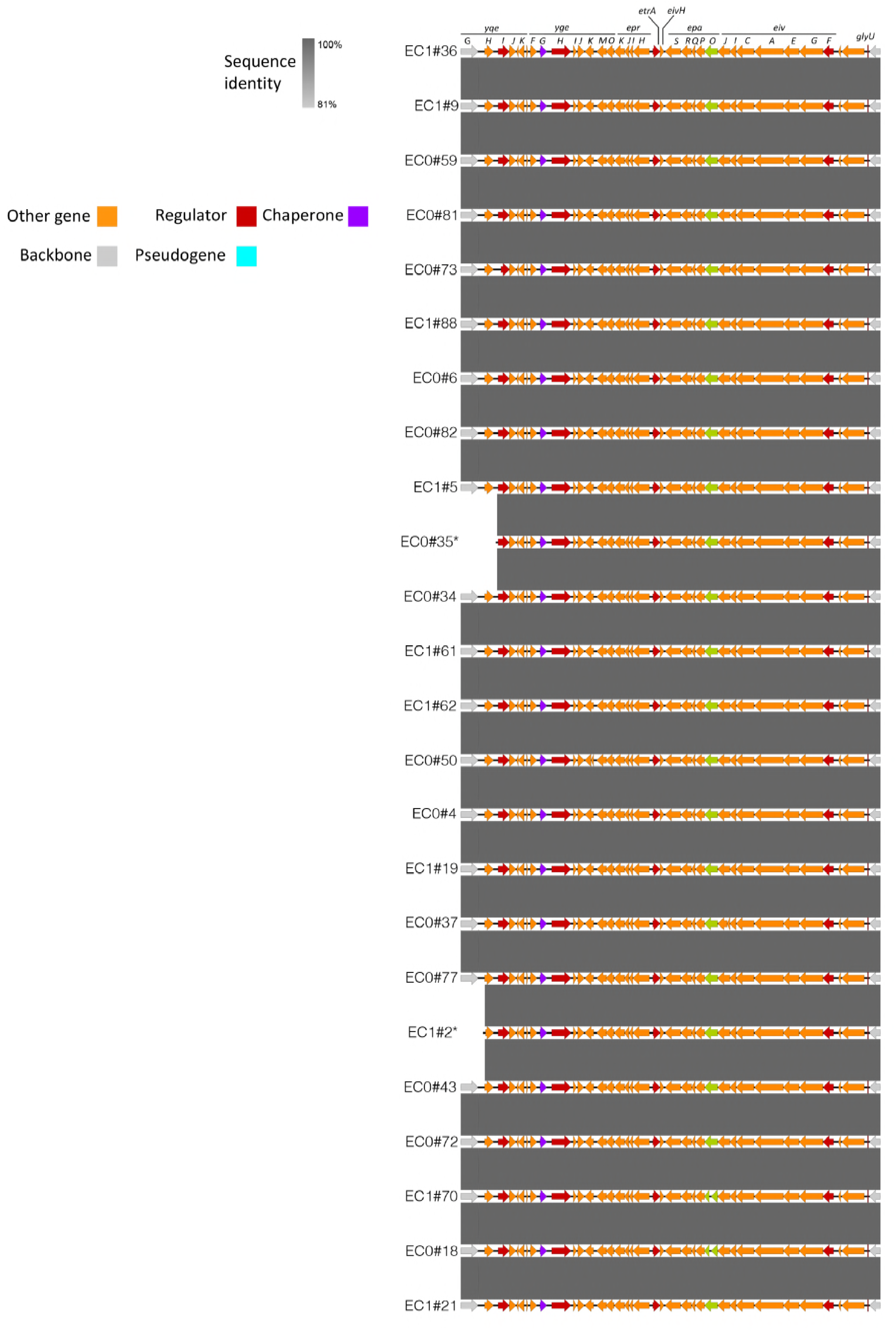
Comparison of the ETT2 operon in 24 ST69 strains. Degree of identity is shown by the level of grey shading as indicated. Genes are colour coded according to putative function as shown.

Next, we analysed other STs within our collection of bacteremic isolates for the presence of the ETT 2 operon (Figure 3). 4 non-ST69 isolates contained an essentially intact ETT2 region, belonging to ST405, 38, 362 and 349. These were group into phylogroups F, D, unknown and unknown respectively; all are closely related to ST69 (supplementary Figure 1). Other strains showed variable loss and/or degradation of the locus as previously described, Notably, none of the common epidemic strain ST131 (phylogroup B2) contains any elements of this ETT2 region – one representative example is shown at the bottom of Figure 3.

**Figure 3.**
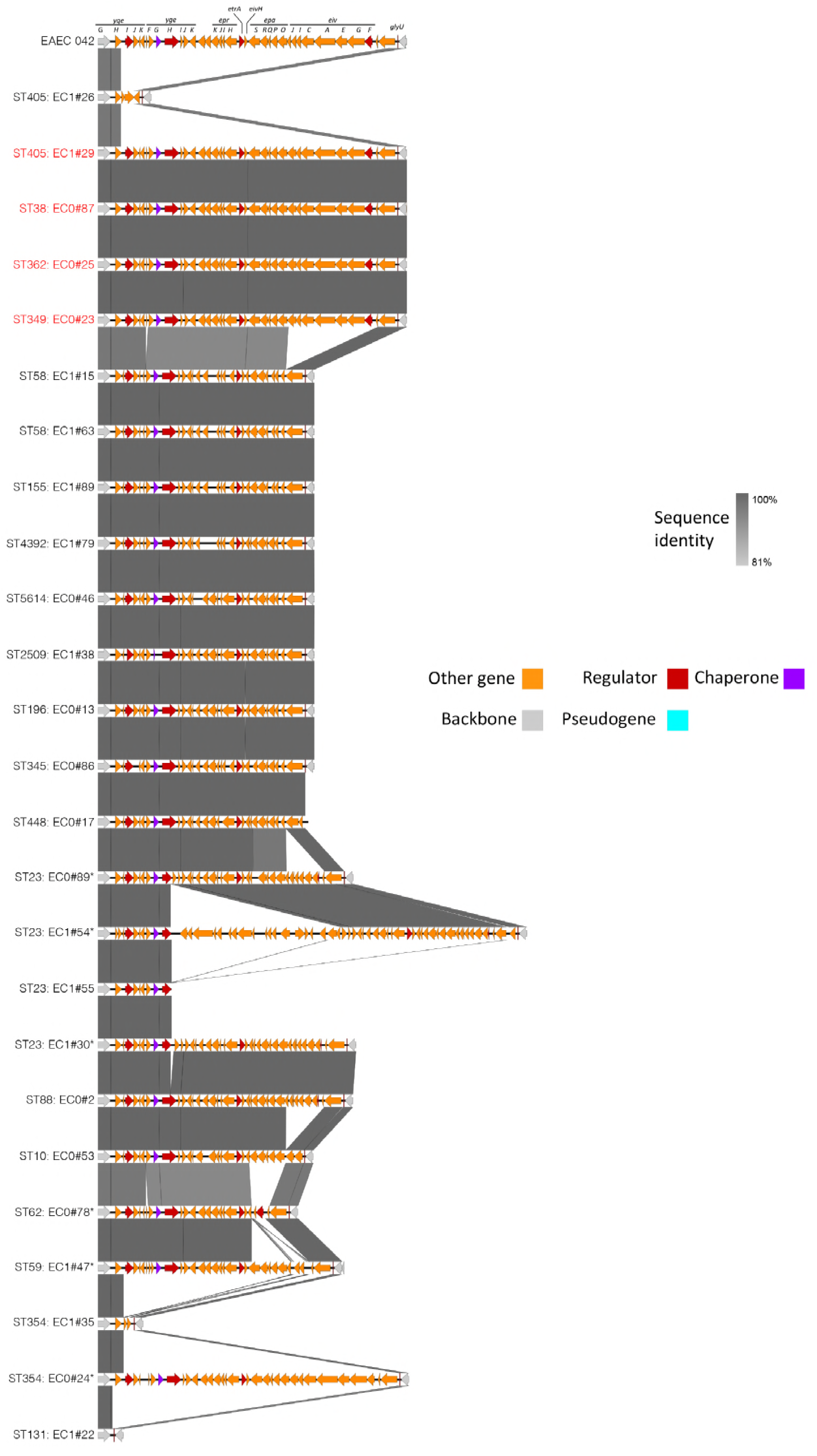
Comparison of elements of the ETT2 operon found in non-ST69 strains. Degree of identity is shown by the level of grey shading as indicated. Genes are colour coded according to putative function as shown. Strains containing an essentially intact ETT2 operon are shown highlighted in red.

Closely associated with an intact ETT2 region is a group of genes related to type III secretion effectors adjacent to the *selC* tRNA gene(12, 15). Two distinct genome insertions were noted at this site: *selC-*A and *selC-*B. Comparison of this region with representative ST69 and other strains compared to EAEC 042 is shown in Figure 4. In EAEC 042 *selC-*A lies between an intact copy of the *selC* gene and a 21 bp direct repeat of the 3’ end of the *selC* tRNA gene. Three backbone genes then intervene (*yicK, yicL, nlpA*) before the region of the *SelC-*B region. *SelC-*A contains mainly phage related genes. *SelC*-B contains homologues of putative type III secretion effectors (*eipB, eipX* and *eipD*), a putative type III effector chaperone, *eicA*, a transcriptional regulator *eilA*, and a gene *eaeX*, which encodes a large protein containing bacterial immunoglobulin repeats with homology to outer membrane adhesion/invasion protein invasion found in *Yersinia* spp. as well as intimins of invasive *E. coli* strains. All ST69 strains in our isolates contained the *SelC-*B locus with over 95% identity to the EAEC 042 region. The variations were found within the central domain of the EaeX product, which contains the bacterial immunoglobulin repeats, with variation in the number of repeats contained within this domain. A similar region was also found in 5 non-ST69 isolates; 4 in ST59 strains and one ST349 strain that also possessed the ETT2 locus. As with the ETT2 locus, the *SelC-*B region was entirely missing in ST131 isolates. The *selC-*A region was largely absent from our isolates but was partially present in one of the ST60 isolates (ECO#72, figure 4).

**Figure 4.**
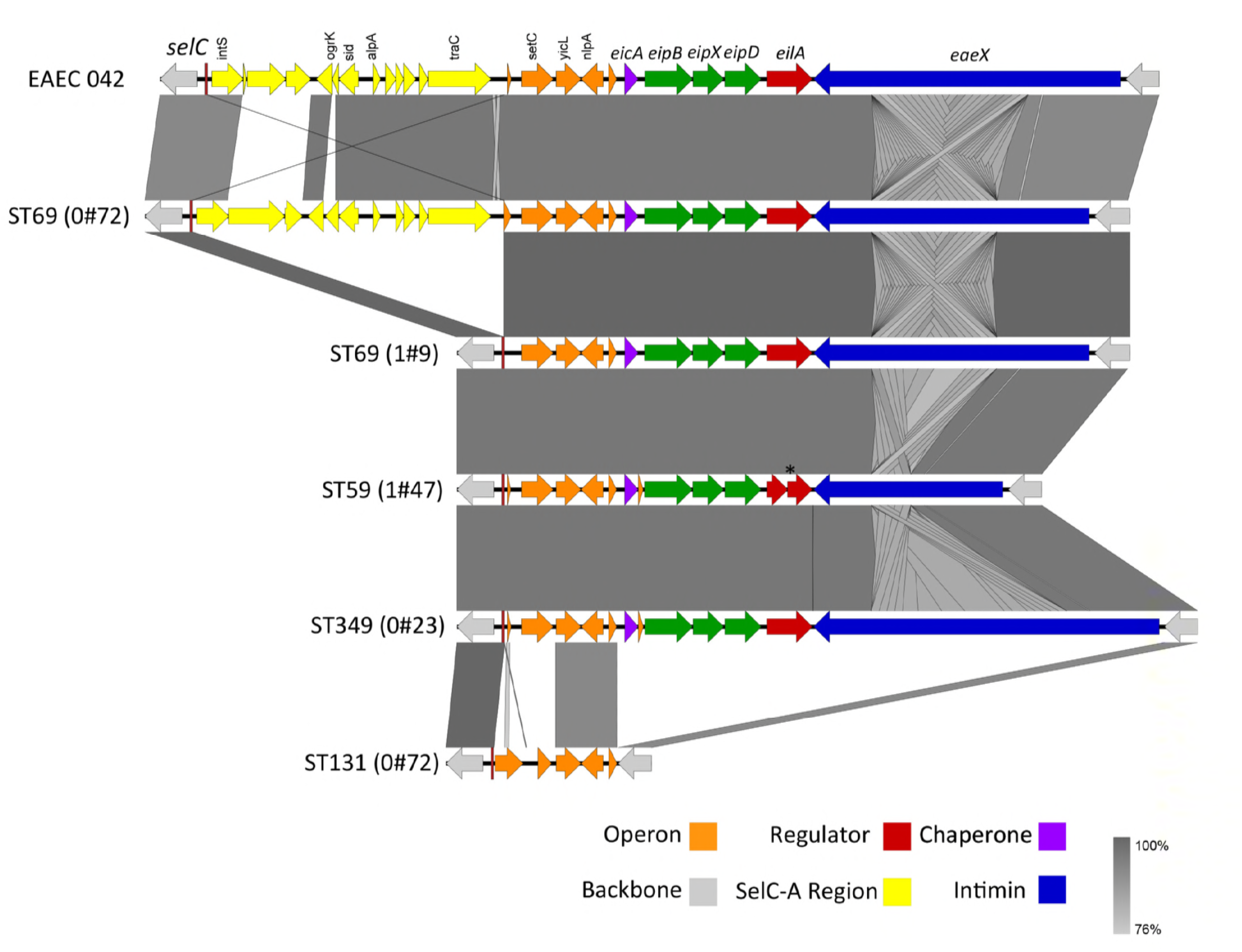
Comparison of the SelC operon in different strains. Degree of identity is shown by the level of grey shading as indicated. Genes are colour coded according to putative function as shown.

*EilA* has been shown to regulate genes within the *SelC*-B region as well as the ETT2 island adjacent to the tRNA *glyU* gene (15). We wished to determine if we could define conditions under which *eilA* was transcriptionally active, and hence activating the ETT2 island. We constructed a reporter gene containing 500 bp of upstream sequence from the *eilA* gene found in the neonatal meningitis associated *E coli* strain CE10(24). Using this reporter in 5 of our ST69 isolates containing the ETT2 locus, we could readily detect reporter gene activity that peaked in the late log phase of growth in equal parts LB and Dulbecco’s Modified Eagle’s Medium (LB:DMEM media) (Fig 5A and B). Previous studies of transcriptional activation of the LEE have shown this is maximal in less rich media designed for growth of eukaryotic cells such as DMEM compared to the rich medium LB(25, 26). Following optimization of growth in different media, we compared transcriptional activity of the *eilA* reporter construct in an ST69 strain grown in LB alone compared to the 1: 1 mixture of LB and DMEM (Fig 5C and D). Growth in the different media was not significantly different but induction of the promoter was much more marked in the LB:DMEM mix. In an attempt to identify proteins potentially secreted into the growth media by ETT2, we compared the pattern of secreted proteins from an ST69 strain with intact ETT2 between the two different media (Fig S2). Two secreted proteins were predominantly found in the bacteria grown in the LB:DMEM mix that we postulated could be potentially secreted by the ETT2. These were cut from the stained gel and subjected to identification by MALDI-MS/MS (Supplementary Table 1). The best matches for these two proteins were the molecular chaperones ClpB and DnaK respectively, which are both molecular chaperones important in refolding aggregated proteins and in protein secretion(27). Neither are putative T3SS substrates.

**Figure 5.**
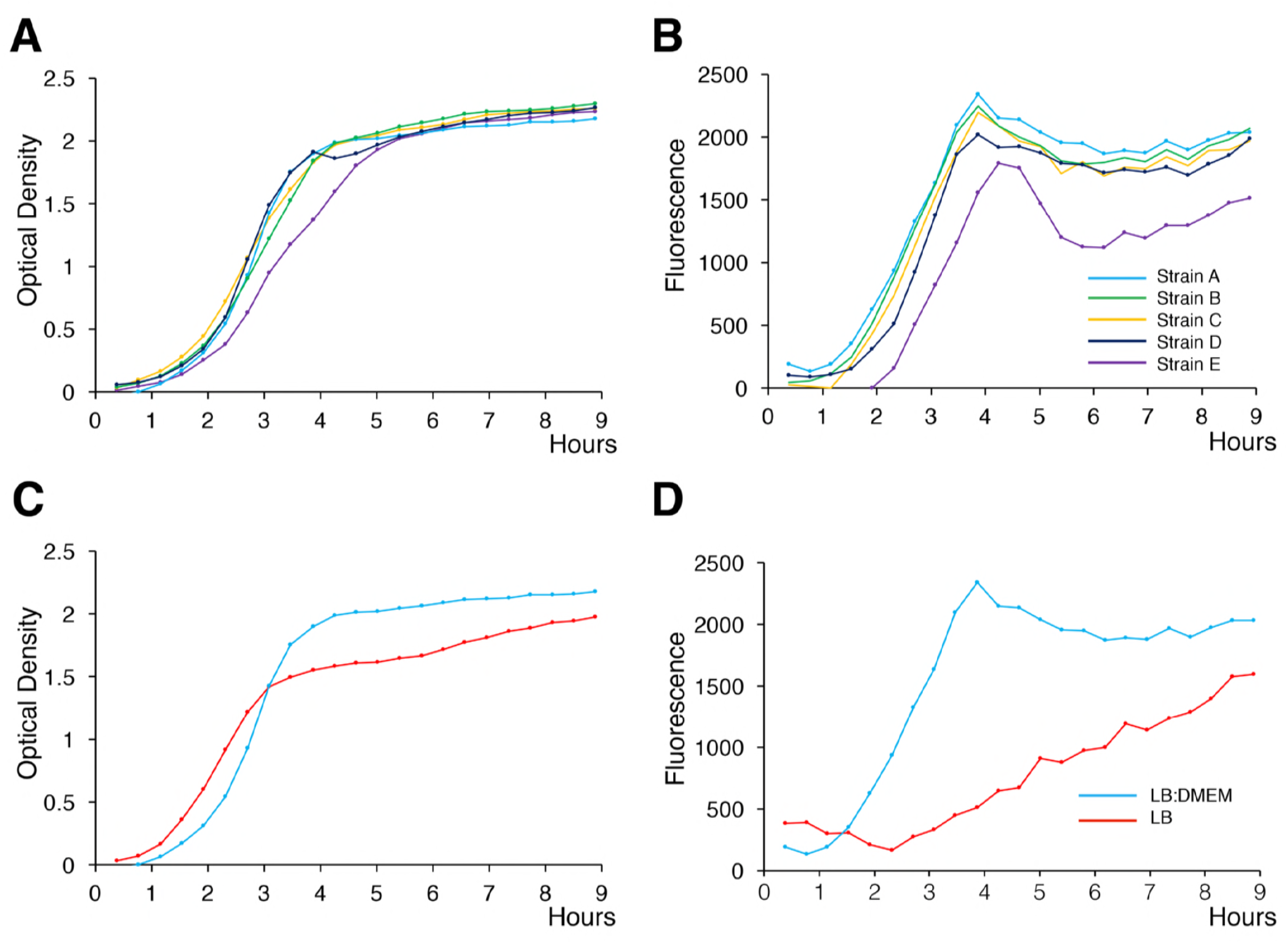
Activity of the eilA reporter in different strains and media. **A** and **B**. Graphs show growth (Optical Density, panels A) and reporter activity (GFP fluorescence, panels B) at the times indicated. The strains are: EC1#2 (A), EC1#19 (B), EC1#5 (C), EC1#21 (D), and EC1#9 (E), all grown in LB:DMEM mixture. Each point is the mean of a triplicate determination; error bars (sem) are contained within the points. **C** and **D** strain EC1#2 is grown in the different media as indicated.

## Discussion

We report here the presence of genomic regions encoding ETT2 and associated putative T3SS effectors within *E. coli* ST69 isolates from bacteremic patients within Scotland. In virtually all of the isolates, the two regions encoding these proteins contained a full complement of genes with no deletion, insertions or inactivating mutations suggesting that the ETT2 and associated effectors could be functionally active. This is in contrast to the vast majority of ETT2 sequences reported to date, which have undergone significant mutational attrition. The conserved nature of the ETT2 sequences reported here strongly suggests that there has been selection pressure for these regions to be conserved within the ST69 lineage.

ST69 belongs to phylogroup D of the *E. coli* lineage. We did not detect ETT2 in *E.coli* of ST131, which is phylogroup B2. Although not completely clear, our data are in agreement with the origin of the different phylogroups as discussed by Ren et al.(12), who suggest that ETT2 is not present in the ancestral B2 phylogroup but was acquired at some point in the evolution of the D group. Subsequent lineages show significant mutational attrition of the ETT2 locus, although our data show strong conservation in the isolates of ST69 we studied here. ST69 is one of the common STs found in bloodstream isolates of *E. coli*. In our collection, ST69 was mostly found in infections acquired from the community(20). The natural environment of these human pathogenic *E. coli* is the gastrointestinal tract; passage into blood is predominantly through ascending infection into the bladder and renal tract. Evolutionary pressure to retain ETT2 might therefore have arisen through its ability to provide a selective advantage in gut colonization and/or in infection of the renal tract.

However, the functional effects of ETT2 remain obscure. In strains with a disrupted ETT2, genetic deletion does seem to confer a changed phenotype, suggesting that even these apparently non-functional regions have a pathogenic role(19). Additionally, experiments in avian strains with ETT2 also suggest a functional role for the ETT2 in pathogenesis(17). ETT2 has also been implicated in the control of gene expression from the locus of enterocyte effacement in enterohemorrhagic *E. coli* O157(28). We could not identify any putative secreted ETT2 substrates from the ST69 strains reported here. A recent study of *E. coli* serotype O2 that causes avian coccobacillosis also failed to identify potential ETT2 secreted proteins, but did find that the intact ETT2 mediated expression and secretion of flagellar proteins, as well as other changes in cell surface behaviour(29). It may be that the conditions under which the ETT2 mediates secretion have not been identified, or that it carries out different functions.

In summary therefore, we show here that the ST69 strain of human pathogenic *E. coli* has an intact genetic locus for ETT2 and associated proteins. The preservation of these sequences in the ST69 strain argue strongly that its functional effects confer a significant selection advantage. However, its exact functional effects remain obscure.

## Materials and Methods

### Sequencing and genome analysis

Whole genome sequencing of 162 strains of E coli from human clinical samples were collected and sequenced as previously described(20). The raw Illumina reads were mapped to the E. coli reference genome EAEC 042 (accession number GCA_000027125.1) using SAMtools mpileup(30) and were called for SNPs through VarScan(31) (read depth ≥2x, variant allele frequency ≥0.08 and p-value ≥0.005). Mobile genetic elements (MGEs) were masked and recombination filtration was performed using Gubbins(32). Maximum likelihood (ML) trees were inferred using RAxML(33) with generalized time-reversible (GTR) model and a Gamma distribution to model site-specific rate variation. One hundred bootstraps were conducted for the support of the SNP based ML phylogenetic tree. Comparison between selected sequences were made and visualised using Easyfig(34)

### Growth and eilA reporter assay

Growth media used in this study were DMEM (Invitrogen, UK), LB, and a 1:1 mix of LB with DMEM. The eilA reporter construct contains a ∼500bp fragment upstream of the eilA promoter from the CE10 strain that was cloned into a plasmid (pAJR70) used in a previous study for the assessment of transcription of ETT1 operons by enhanced green fluorescent protein (GFP) monitoring from liquid culture (35). The plasmid was transfected into the different bacterial strains using standard methods. Chloramphenicol (25µg/ml) was added to media when required for the selection of strains containing the eilA reporter. Induction of GFP in the different media at 37°C was measured using a fluorescence plate-reader (FLUOstar Optima; BMG; Labtech, UK). Optical densities and fluorescence were recorded every 24 minutes for 9 hours.

### Secretion Assay

Secreted protein assays were extracted by trichloroacetic acid precipitation performed as previously described (36). Briefly, overnight LB cultures were diluted 1/100 in 50ml of the culture media and grown for 9 hours before precipitation of secreted proteins. Secreted proteins were resuspended in 150µl of loading buffer and analysed by SDS-PAGE.

### Accession numbers

Illumina sequences are deposited in the European Nucleotide Archive (ENA: www.ebi.ac.uk/ena) under project PRJEB12513.

## Acknowledgements

The work was funded by the Scottish Executive via the Chief Scientists Office through the provision of a grant to establish the Scottish Healthcare Associated Infection Prevention Institute (SHAIPI). The funders had no role in the study design, data collection and interpretation, or the decision to submit the work for publication.

---

**Figure S1. Maximum likelihood phylogenetic tree of the strains shown in Figure 3.** Strains are colour coded according to their ST as shown. The 4 non-ST69 strains with an essentially intact ETT2 operon are indicated by an asterisk.

**Figure S2. Secreted proteins from ST69 grown in different media.** Secreted proteins from the ST69 strain grown in the media indicated were analysed by SDS-PAGE. Molecular weight markers (M) in kDa are shown to the left of the gel. The two proteins that were further analysed by tandem mass spectrometry are shown boxed and labelled 1 and 2.

**Supplementary Table 1. MASCOT summary data for the excised bands from Figure 6.** The summary header gives the top matching GenBank protein database accession number, the relative molecular mass, the total score for the matched protein, the number of peptides matched (Matches), unique peptides matched (Sequences), and exponentially modified protein abundance index (emPAI). The table columns show the identification number of the peptide (Query), the observed mass/charge ratio (Observed), the observed relative molecular mass (Mr (expt)), the calculated relative molecular mass (Mr (calc)), the difference in parts per million between the observed and calculated masses (ppm), number of missed cleavage sites (Miss), the score for each peptide which is -log10(Expect), the probability of observing the peptide by chance (Expect), the rank of the peptide match (Rank), the identity of the peptide match (Unique), where U signifies the peptide is unique to one protein family member, and the peptide sequence using the one letter amino-acid code (Peptide), where the residues that bracket the peptide sequence in the protein are also shown, delimited by periods.

